# Extraordinarily wide genomic impact of a selective sweep associated with the evolution of sex ratio distorter suppression

**DOI:** 10.1101/006981

**Authors:** EA Hornett, B Moran, LA Reynolds, S Charlat, S Tazzyman, N Wedell, CD Jiggins, GDD Hurst

## Abstract

Symbionts that distort their host’s sex ratio by favouring the production and survival of females are common in arthropods. Their presence produces intense Fisherian selection to return the sex ratio to parity, typified by the rapid spread of host ‘suppressor’ loci that restore male survival/development. In this study, we investigated the genomic impact of a selective event of this kind in the butterfly *Hypolimnas bolina*. Through linkage mapping we first identified a genomic region that was necessary for males to survive *Wolbachia*-induced killing. We then investigated the genomic impact of the rapid spread of suppression that converted the Samoan population of this butterfly from a 100:1 female-biased sex ratio in 2001, to a 1:1 sex ratio by 2006. Models of this process revealed the potential for a chromosome-wide selective sweep. To measure the impact directly, the pattern of genetic variation before and after the episode of selection was compared. Significant changes in allele frequencies were observed over a 25cM region surrounding the suppressor locus, alongside generation of linkage disequilibrium. The presence of novel allelic variants in 2006 suggests that the suppressor was introduced *via* immigration rather than through *de novo* mutation. In addition, further sampling in 2010 indicated that many of the introduced variants were lost or had reduced in frequency since 2006. We hypothesise that this loss may have resulted from a period of purifying selection - removing deleterious material that introgressed during the initial sweep. Our observations of the impact of suppression of sex ratio distorting activity reveal an extraordinarily wide genomic imprint, reflecting its status as one of the strongest selective forces in nature.

## Author summary

The sex ratio produced by an individual can be an evolutionary battleground. In many arthropod species, maternally inherited microbes selectively kill male hosts, and the host may then in turn evolve strategies to restore the production or survival of males. When males are rare, the intensity of selection on the host may be extreme. We recently observed one such episode, in which the population sex ratio of the butterfly *Hypolimnas bolina* shifted from 100 females per male to near parity through evolution of a suppressor gene. In our current study, we investigate the hypothesis that the strength of selection was so strong in this case that the genomic impact would go well beyond the suppressor gene itself. Following location of the suppressor within the genome of *H. bolina*, we examined changes in genetic variation at sites on the same chromosome as the suppressor. We show that an extraordinarily wide area of the genome was affected by the spread of the suppressor. Our data also suggest that the selection may have been sufficiently strong to introduce deleterious material to the population that was later purged by selection.

## Introduction

In 1930, Fisher noted that the strength of selection on the sex ratio was frequency dependent [1]. As a well-mixed outbreeding population progressively deviates from a 1:1 sex ratio, selection on individuals to restore the sex ratio to parity becomes stronger. In natural populations, a principle cause of population sex ratio skew is the presence of sex ratio distorting elements, in the form of either sex chromosome meiotic drive [2], or cytoplasmic symbionts [3]. In some cases, these elements can reach very high prevalence, distorting population sex ratios to as much as 100 females per male [4], and producing intense selection for restoration of individual sex ratio to 1 female per male. The most common consequence of this selection pressure is the evolution of systems of suppression – host genetic variants that prevent the sex ratio distorting activity from occurring. Suppressor genes are known for a wide range of cytoplasmic symbionts and meiotic drive elements [2,5,6].

The evolution of suppression of *Wolbachia* induced male-killing activity in the butterfly *Hypolimnas bolina* represents a compelling observation of intense natural selection in the wild. Female *H. bolina* can carry a maternally inherited *Wolbachia* symbiont, wBol1, which kills male hosts as embryos [7]. The species also carries an uncharacterised dominant zygotically acting suppression system that allows males to survive infection [5]. Written records and analysis of museum specimens indicate this symbiont was historically present, and active as a male-killer, across much of the species range, from Hong Kong and Borneo through to Fiji, Samoa and parts of French Polynesia [8]. Evidence from museum specimens also indicates that host suppression of male-killing had a very restricted incidence in the late 19^th^ century, with infected male hosts (the hallmark of suppression) being found in the Philippines but not in other localities tested. By the late 20^th^ Century, suppression of male-killing was found throughout SE Asia, but not in Polynesian populations where the male-killing phenotype remained active [9]. The most extreme population was that of Samoa, where 99% of female *H. bolina* were infected with male-killing *Wolbachia*, resulting in a population sex ratio of around 100 females per male [4]. However, following over 100 years of stasis on Samoa, rapid spread of suppression of male-killing activity of the bacterium was finally observed between 2001 and 2006, restoring both individual and population sex ratio to parity [10].

When selection occurs at a locus, it is expected to leave a genomic imprint beyond the target of selection, as a result of genetic hitch-hiking. During the spread of a beneficial mutation, any variant that is initially associated with the selected locus through linkage will also increase in frequency [11]. When selection is strong, such selective sweeps may increase the frequency of linked variants across a broad genomic region [12]. Importantly, the extent of a sweep will most depend on the selection pressure in the first few generations, before recombination has broken down associations between the target of selection and linked variants. Where sex ratio distorters are common, the selection pressure in these first generations may be very strong indeed. It is thus likely that selection on the sex ratio will influence linked material over an extraordinarily broad genomic region, as compared to many other selective regimes. That is, the episode of selection is likely to have a very wide genomic impact.

In this paper, we first mapped a genomic region in SE Asian butterflies that was required for male survival in the presence of *Wolbachia*. We then investigated the impact of the recent spread of the suppressor in Samoa on the pattern of variation around this region. To this end, we initially developed theory to predict the impact of suppressor spread on linked genetic variation. We then directly observed changes in the frequency of genetic variants surrounding the suppressor locus by comparing the pattern of genetic variation in *H. bolina* specimens collected in Samoa before (2001) and after the sweep (2006 and 2010). By examining post-sweep samples at two time points we were additionally able to track allele frequency changes following the initial sweep. The data revealed the broadest sweep recorded to date in nature. We further suggest that the suppressor was likely derived through immigration, and that the sweep may have introduced deleterious material subsequently subject to purifying selection.

## Results and Discussion

### Location of a region required for male survival in the H. bolina genome

*Hypolimnas bolina* has 31 chromosomes and a total genome size of 435 MB [13] [14]. Genetic markers spanning the genome were developed using a targeted gene approach informed by conservation of synteny in Lepidoptera, with the sequence of *H. bolina* orthologs obtained through Roche 454 transcriptome sequencing (see Methods and Materials). These markers were then tested for co-segregation with suppression in order to identify the linkage groups associated with male host survival. Female butterflies from SE Asia that carried both the *Wolbachia* and the suppressor allele, were crossed with males from the French Polynesian island Moorea (where suppression is absent). These resulting F1 daughters (who inherited *Wolbachia* from their SE Asian mother) were then backcrossed to Moorea males to create a female-informative family for identification of loci linked to the suppressor. The absence of recombination in female Lepidoptera means a SE Asia allele necessary for male survival that is present in the F1 mother will be present in all her sons (as if they lack it, they die), but show normal 1:1 segregation in her daughters (Fig. 1). From 8 initial genomic loci screened, one locus orthologous to sequence on chromosome 25 in the moth *Bombyx mori* showed this pattern of inheritance whereby all 16 sons carried the same maternal allele of SE Asia origin while 8 daughters showed Mendelian segregation at this locus (probability of observing this pattern of segregation in sons = (1/2)^16^: p<0.0001). We then obtained a further 11 markers in this linkage group. Candidates were identified initially via synteny to *B. mori*, and then confirmed as showing co-segregation with the original marker and as being associated with male survival, in the female-informative family. In this way, a suite of 12 suppressor-linked markers (A-L) were developed, all of which followed the presumed pattern of inheritance of the suppressor - that of presence in all 16 sons and half daughters.

**Figure 1.**
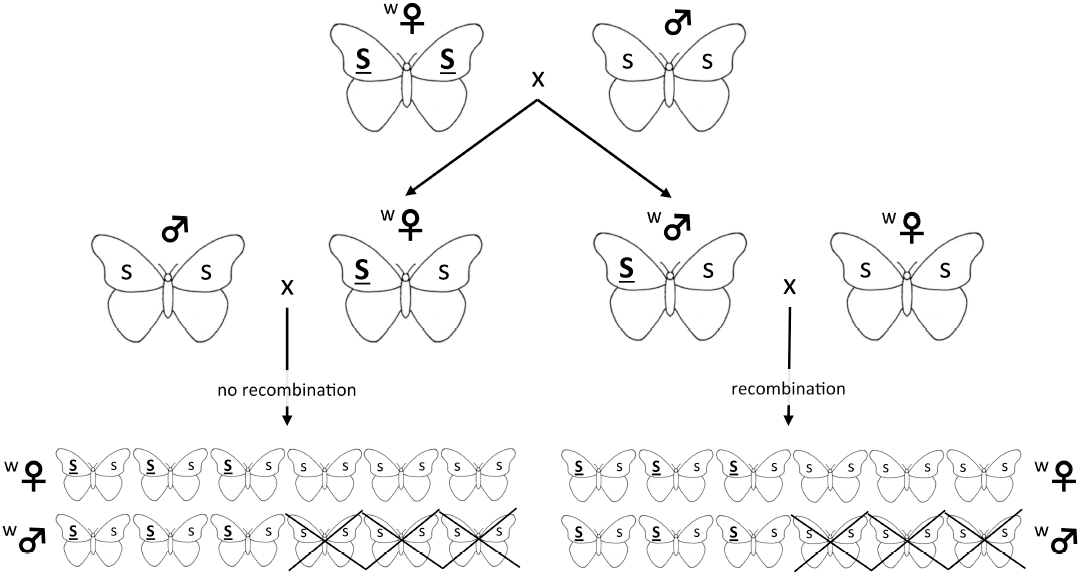
Mapping of the *Hypolimnas bolina* genomic region surrounding the suppressor of male-killing. A *Wolbachia* infected (denoted by ‘w’) female that was homozygous for the suppressor allele (***<u>SS</u>***) was crossed to a uninfected male that did not carry this allele (*ss*). To produce a female-informative family, *Wolbachia*-infected heterozygous daughters (***<u>S</u>****s*) from this pairing were in turn crossed to uninfected males lacking the suppressor. Because there is no recombination in female Lepidoptera, male survival is associated with inheritance of the linkage group carrying the suppressor, and suppressor-linked loci can be identified as those present in all surviving F2 sons (those marked with a cross die) but only 50% of F2 daughters. To produce a male-informative family, *Wolbachia*-infected heterozygous sons (***<u>S</u>****s*) from the original parental cross were crossed to infected females lacking the suppressor. Using this cross, members of the suppressor-associated linkage group were mapped relative to each other through the pattern of recombination in the F2 daughters. The location of the suppressor was ascertained as the genomic region that was present in all surviving F2 sons.

A linkage map for this chromosome was then constructed, with the region within it required for male survival identified by the exclusion of recombinants. This was achieved by examining segregation of alleles from sons of the SE Asia x Moorea cross above that were mated to *Wolbachia*-infected Moorea (non-suppressor) females (creating a male-informative family). Over 300 recombinant daughters were obtained that were used to create a linkage map of the 12 suppressor-linked markers. These markers covered a 41cM recombination distance and were syntenic with *B. mori* (Fig. 2). The suppressor locus was localized to a region within this chromosome by excluding linked loci where the SE Asia derived paternal allele was absent in one or more sons (indicating the genomic region containing the SE Asia allele was not necessary for male survival). Three suppressor-linked alleles, all in the +11 to +12 region, were retained in all 60 sons, whereas the 9 markers proximal and distal to these were excluded by the presence of one or more recombinants (Fig. 2). Binomial sampling rejected the null hypothesis of no association between the +11/+12 genomic region and male survival (p=(1/2)^60^). Thus we posit that the suppressor lies between marker C at +8 (excluded by one recombinant) and marker G at +17 (excluded by two recombinants).

**Figure 2.**
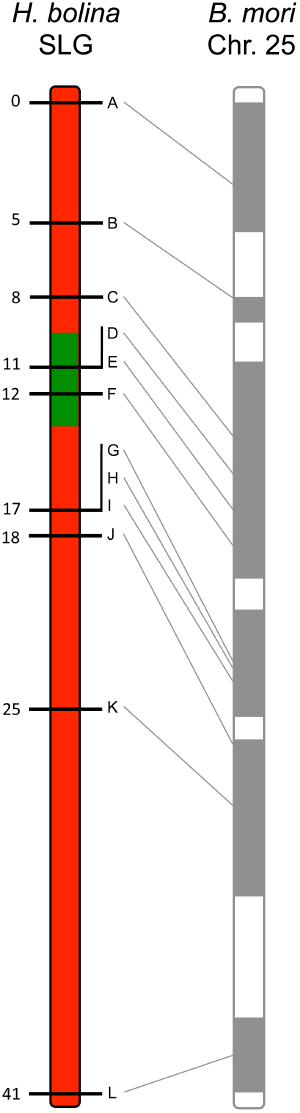
Recombinational map of the *H. bolina* chromosome carrying the suppressor locus. Depiction of the suppressor linkage group (SLG), with the 12 linked markers A-L. The genomic region containing the suppressor locus is highlighted in green. Numbers represent distance from the distal marker in cM as estimated using JoinMap. The broadly syntenic *B. mori* chromosome 25 is given for reference.

### The impact of suppressor spread on linked genetic variation in Samoa

#### a) Modelling a selective sweep driven by suppressor spread

Models of selective sweep dynamics commonly utilize a fixed selective coefficient. In contrast, the intensity of selection is dynamic in our system, as the benefit of rescuing a male relates to population sex ratio, which depends on the frequency of male-killing *Wolbachia* and the frequency of suppression: as the suppressor spreads, the population sex ratio shifts towards 1:1 thus reducing the selective pressure for further spread. We previously modelled the spread of the suppressor in this system using deterministic simulation [15]. This model tracks genotype frequency in males and females separately. For each sex, zygotes are formed by random union of male and female gametes, creating a genotype frequency distribution in zygotes. Male-killing then creates selection between zygotic and adult phases, with any *Wolbachia*-infected males lacking the *S* suppressor gene (i.e. those with genotype *ss*) being killed shortly following formation (early in development). Thus the adult pool of male genotypes differs from the zygotic pool, and has a higher frequency of suppression than the adult female pool. Fisherian selection is then implicit in the model, with the suppressor increasing in frequency between the zygote and adult phase by virtue of being overrepresented in male hosts that contribute 50% of the genetic material to the next generation.

In the Samoan population of *H. bolina*, *Wolbachia* was initially at 99% prevalence in females [4], creating an extremely biased population sex ratio. Under such circumstances a novel suppressor mutation will rise to 50% frequency in adult males in the first generation (all surviving F1 males are heterozygous *Ss*), and to 25% frequency in the subsequent set of zygotes. Males that survive *Wolbachia* infection by carrying the suppressor are now induced to express another reproductive manipulation commonly employed by *Wolbachia* - cytoplasmic incompatibility (CI) against uninfected females [16]. During CI, the progeny of uninfected females sired by infected males die during embryogenesis, such that the presence of infected males (generated through suppressor action) reduces the fitness of uninfected females relative to infected females (who experience no such reduction in offspring viability following mating with infected males). Despite losing its male-killing ability, the induction of CI through infected males allows *Wolbachia* to remain at or near fixation. In consequence selection on the suppressor is also maintained. Indeed fixation for *Wolbachia* in males and females is observed after the spread of the suppressor is present [10].

The expected dynamics of the suppressor locus in this system is given in Figure 3a, following the trajectory given for *r* = 0 (zero recombination with the suppressor). We elaborated the model of suppressor spread to quantify expected effects on linked loci under the assumption that the suppressor and linked alleles were cost-free, and that spread was occurring through an unstructured, panmictic population (see Text S1 for full details). We then modified this model to examine the effect of suppressor spread on levels of association (linkage disequilibrium) between pairs of linked loci, each with two alleles. Within the model, the suppressor locus can have the wild type allele *s*, or the male-killing suppression allele *S*. The second locus is linked to the suppressor locus and has two selectively neutral alternative alleles denoted *A* and *a*. The model tracks the change in gametic frequencies from one generation to the next. There are therefore four different basic gamete types: *AS*, *As*, *aS*, and *as*. Our individuals are diploid, so these four basic gamete types give nine possible basic individual genotypes: *AASS*, *AaSS*, *aaSS*, *AASs*, *AaSs*, *aaSs*, *AAss*, *Aass*, and *aass*. This is further complicated by the need to record whether individuals are infected or uninfected. Since infection is maternally inherited we can count infection status as part of the genotype of an individual, giving us a total of eighteen genotypes (the nine above, with each having infected or uninfected status).

**Figure 3.**
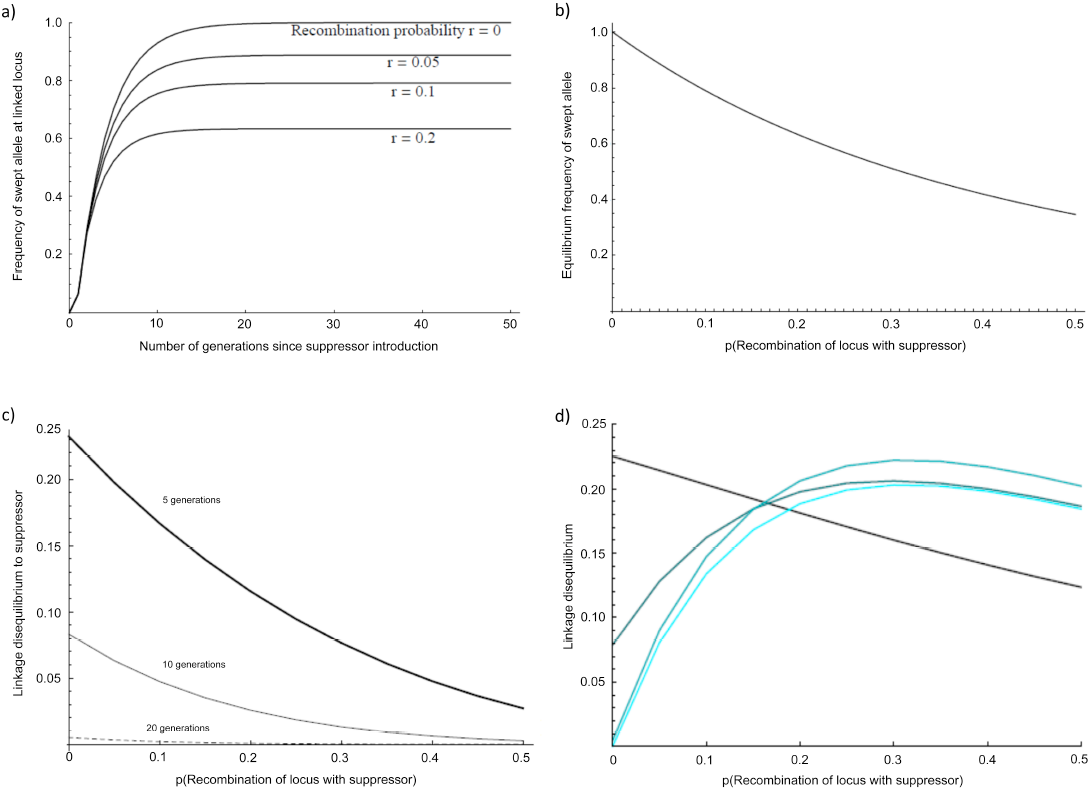
Predicted impact of selective sweep on the frequency and association of linked variants. 3a) Change in the frequency of initially associated linked variants over time at different recombination distances, 3b) Equilibrium frequency of initially associated linked variants at different recombination distances, 3c) Linkage disequilibrium between loci at different recombination distance from suppressor after 5, 10 and 20 generations of selection, 3d) Local linkage disequilibrium between alleles with recombination probability of 0.01 at different recombination distances from the suppressor. The four curves show the situation after 5, 10, 20, and 40 generations (increasing lightness of colour corresponding to greater number of generations).

Given a set of genotype frequencies for males and females, we can derive the expected gametic frequencies (incorporating recombination in males), and assuming random mating we can calculate the frequencies of offspring genotypes in the next generation. We can then apply selection: we assume that the infection kills all males lacking the suppressor allele (*ss*), half of the males that are heterozygous (*Ss*) for the suppressor, and none of the males that are homozygous (*SS*) for the suppressor. This partially dominant rescue mirrors patterns of male survival in our mapping crosses, in which about half of Ss males survive. For simplicity we assume no other selective effects. This process allows us to track both gametic and genotypic frequencies through time.

We iterated this process for the Samoan situation where 99% of females are infected with a fully penetrant male-killer, in which males only survive in the presence of the suppressor allele, and supposing that the suppressor gene initially appeared on a single infected immigrant male of genotype *AASS*. The selected locus (*r* = 0) is observed to come to equilibrium within 10-15 generations, which is consistent with the observed rapid spread of the suppressor (Fig. 3a). Overall, our panmictic model predicts the sweep would cover the entire suppressor-containing chromosome (Fig. 3b), with markers at recombination distance 0.5 to the suppressor locus being swept to 35% frequency. The cause of this breadth is selection in the first generation, in which effectively all males in the mating pool are heterozygous for the chromosome carrying the suppressor (thus this chromosome rises to an initial frequency approaching 0.25).

We further examined how the sweep would impact on the level of linkage disequilibrium (LD) within the linkage group, both in terms of LD to the suppressor locus (Fig. 3c), and local LD (disequilibrium between pairs of loci other than the suppressor) (Fig. 3d; Text S1). After 10 generations of selection, association between the suppressor locus and linked loci was weak, and at 20 generations it was expected to be undetectable for a sample of 50 individuals. In contrast, local LD rose and fell over a longer time period. Local LD was generated by the sweep initially taking locally associated variants in concert. Where the loci are closely linked to each other, the LD generated was expected to be retained even by 40 generations. The extent of LD was maximized at intermediate recombination distances from the suppressor locus, where swept alleles were at intermediate frequency when at equilibrium.

#### b) Observations of the effect of suppressor spread on patterns of genomic variation between 2001 and 2006

We investigated the change in frequency of genetic variants associated with the spread of suppression of male-killing in *H. bolina* on the island of Upolu, Samoa, through comparison of allelic profiles before and after the spread of suppressor. To this end, intronic regions of the 12 suppressor-linked markers were sequenced for butterflies from before (2001, n=48), and after (2006, n=48) the spread of the suppressor. Allele frequency in each sample was estimated using PHASE and the results checked manually. The history of selection was examined directly by testing for differences in allelic frequency distributions at each loci between the pre- and post-sweep samples and by estimating Fst between these samples as a standardized metric of change, and indirectly by the estimating LD between loci - a hallmark of recent selection. Changes were compared to a control group of 9 unlinked loci located throughout the genome that would indicate the presence of genome-wide effects, as opposed to suppressor linkage group specific, occurring in this period.

Significant changes in the allele frequency distribution between the 2001 and 2006 population samples were observed at 11 of the 12 markers along the chromosome previously identified as carrying the suppressor in SE Asian butterflies (Fig. 4; all 11 with p<0.01 after sequential Bonferroni correction: Fig. S1). Only the most distant marker from the suppressor (marker L) showed no evidence of change. Overall, significant deviations were observed over a 25cM region surrounding the location of the suppressor previously mapped in SE Asian butterflies. In contrast, none of the 9 un-linked markers showed evidence of a change in allelic frequency (NS at p=0.05 after sequential Bonferroni correction; Fig. S2), allowing us to reject demographic factors and drift as a cause of changes in allele frequency in the suppressor linkage group.

**Figure 4.**
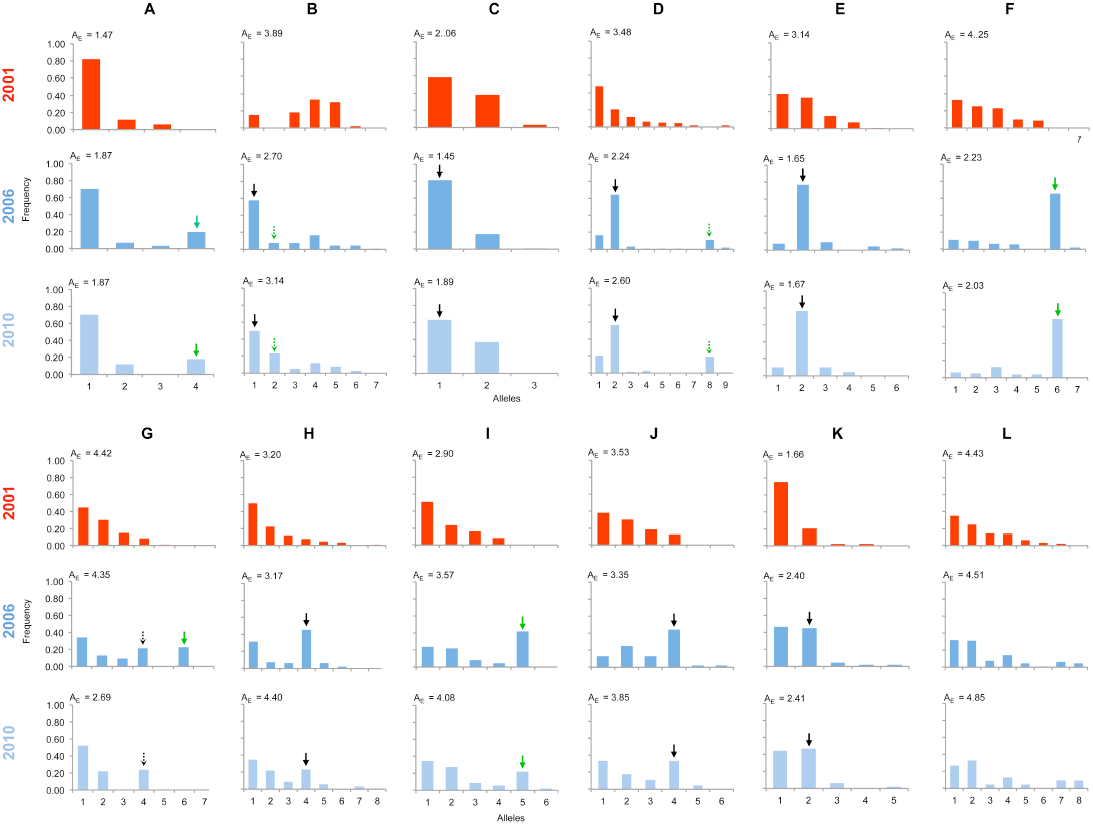
Allelic frequency distributions across the linkage group carrying the suppressor in 2001, 2006 & 2010. The major allelic variant for each locus (A-L) that has increased in frequency during the sweep is indicated with a solid arrow, while the secondary allele that increases (in 3 cases) is indicated with a dotted arrow. Where the allele is novel in 2006/2010 the arrow is green. Results of statistical comparison presented in Fig. S1. Effective allele numbers (A_E_) are indicated on each graph.

For each of the suppressor-linked loci that showed significant change from 2001-2006 we identified the allele that had ‘swept’ alongside the suppressor. An expectation for a selective sweep is that allelic distributions will be disturbed by an increase in frequency of one (or sometimes two) alleles initially associated with the target of selection, which would be paralleled in a co-ordinated decline in frequency of the other alleles. Because decline is spread across multiple alleles, the expectation is that a swept allele should be the greatest contributor to heterogeneity between allelic frequency distributions, and thus identifiable by the largest standardized residual in heterogeneity tests. We performed this analysis of residuals at 10 of the loci where heterogeneity between samples was observed (one locus was not suitable for analysis in this manner, as there were only two allelic variants, thus by definition each allele contributes equally to heterogeneity). At each of the 10 loci, a single allele increasing in frequency from 2001 to 2006 contributed the largest residual to heterogeneity analysis, and thus can be regarded as the swept allele. However in 3 cases (loci B, D & G), removal of this allele did not restore homogeneity, and a second allele that increases in frequency can additionally be identified as contributing to the heterogeneity observed, albeit with a lower magnitude of frequency change (swept alleles annotated with arrows in Fig. 4).

We also calculated the Fst statistic for each locus as a standardized magnitude of difference between the 2001 and 2006 population samples (Fig. 5). Fst differentiation was highest in the 5cM region in which the suppressor had been previously located in SE Asian butterflies (Fst=0.2-0.3), and lower, but significant, differentiation was found over the 25cM region surrounding the locus (p<0.01 at all loci after sequential Bonferroni correction), making this selective sweep the broadest observed to date in nature. Fst differentiation was either absent or present at a very low level (Fst<0.03) in the group of 9 unlinked markers (Fig. S3).

**Figure 5.**
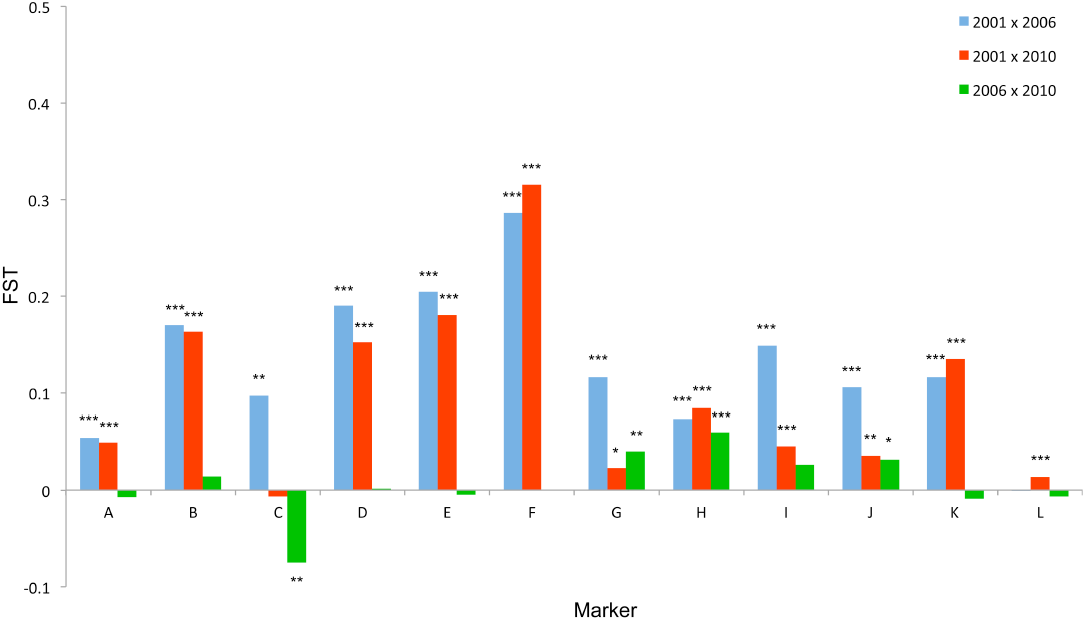
Fst standardized population genetic differentiation between samples from different time points for each locus along the chromosome carrying the suppressor. Blue: 2001-2006; Red: 2001-2010; Green: 2006-2010. Statistically significant deviations from Fst=0 given by *** (p<0.001), ** (p<0.01) * (p<0.05) uncorrected for multiple tests.

The pattern of LD also reflected an episode of selection in this genomic region. Our model above predicted that LD between the swept loci and the target of selection (global LD) would exist only in the very early phases of the sweep (10-20 generations), but that the sweep would create multiple local associations between closely linked loci that would be retained over longer periods (local LD) (Fig. 3c, 3d). In accord with this, there was little evidence of LD between loci in the 2001 pre-suppressor sweep sample, but strong associations between variants at closely linked loci (r=0.01-0.02) in the 2006 sample (Fig. 6).

**Figure 6.**
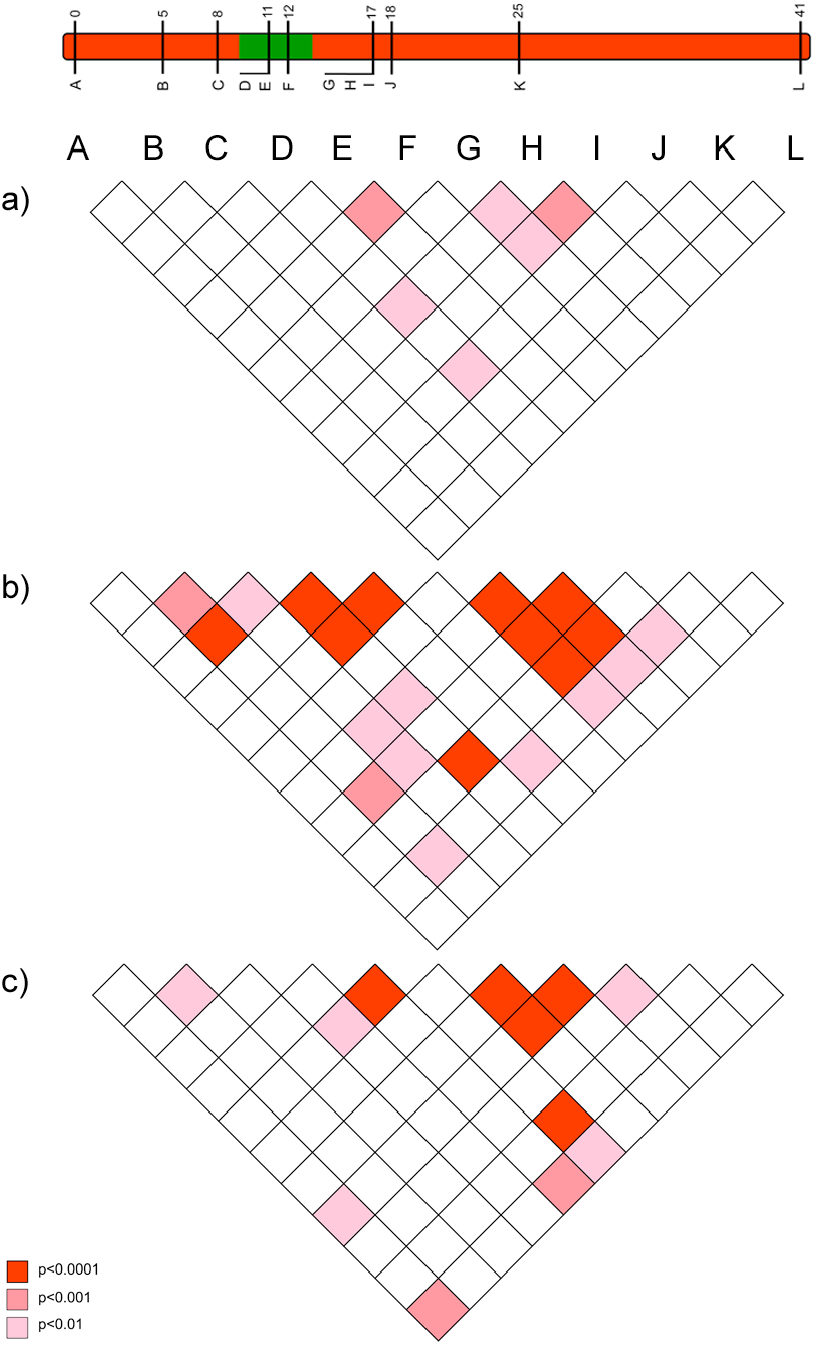
The pattern of linkage disequilibrium (association) between loci observed in 2001 (pre-sweep), 2006 (immediately post-sweep) & 2010 (4 years post sweep) population samples. Pattern of linkage disequilibrium (LD) between suppressor-linked loci (A-L) for a) 2001 (pre-sweep), b) 2006 (immediately post-sweep), and c) 2010 (4 years post-sweep). Significance denoted by colour: deep red – significant LD as measured at p<0.001; mid red – significant LD as measured at p<0.01; pink – significant LD as measured at p<0.05, all uncorrected for multiple tests. Depiction of suppressor linkage group given for reference.

Our data indicate that the genomic region identified in SE Asian butterflies as carrying a suppressor locus was also the focus of selection in the selective sweep in Samoa. This observation has two possible explanations. First, the suppressor mutation may have been introduced into Samoa by migration. Given the suppressor is absent in the nearest island groups, American Samoa and Fiji, suppressor introduction would be associated with a long distant migrant. Second, the genomic region may represent a hotspot for suppressor mutation, derived independently in Samoa by *de novo* mutation.

The presence of novel swept alleles at linked loci indicates that migration is the most parsimonious explanation for suppressor origin. Variants not present in the 2001 sample were observed to be the main ‘swept’ allele at 4 of the 11 loci at which significant change was detected (indicated with green arrows in Fig. 4). At two of these loci (A & I), the invading allele was defined by a SNP being not present in the 2001 sample, whereas the other two alleles represented different combinations of existing SNPs. The four loci were in three genomic locations spaced over 17cM and showed no evidence of linkage disequilbrium (association) in the 2001 pre-sweep sample, and thus they can be treated as independent (Fig. 6). We believe it is very unlikely that all four alleles existed in the 2001 population but were not sampled. We reason that derivation of alleles selected by correlation to a *de novo* suppressor mutation would occur in proportion to their frequency in the pre-sweep population. There is an upper confidence limit for the frequency of alleles absent in the 2001 sample of 0.0307 (binomial sampling n=96 alleles, 95% confidence interval). Taking the conservative assumption that each absent allele existed at the upper confidence level for frequency, four or more unsampled alleles being present in a sample of 11 as targets of selection occurs with a chance of p=0.00025 (calculated from binomial sampling distribution: 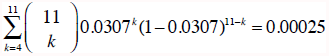, where each locus has a chance of 0.0307 of having a previously unsampled allele, and obtaining four or more such alleles from 11).

Thus the presence of four novel linked variants argues against a model of *de novo* mutation to suppression. In addition, two of the three secondary ‘swept’ alleles (those that contributed significantly to locus heterogeneity between the 2001 and 2006 samples, after the main swept allele had been removed) were also novel in 2006 (markers B & D - denoted by dotted green arrows in Fig. 4). We thus conclude a migratory source for the suppressor represents a more parsimonious explanation for the origin of suppression, although definitive proof of this awaits fine-scale mapping and characterization of the suppressor mutation itself both within Samoa and from possible source populations (e.g. SE Asia).

Notwithstanding the broad sweep observed, changes across the suppressor chromosome were not as profound as our model predicted. It is likely that the assumption of panmictic population structure within the model accounts for this difference. In natural populations, the suppressor mutation will occur initially at one geographical location, and the mutation would then spread away from this location each generation in a wave of advance. This spatial spread of suppression was observed in Savai’i, the neighbouring island to Upolu, and took at least one year (10 generations) to complete [10]. Each generation of spatial advance would have seen a less broad local sweep, as association of loci to the suppressor locus would have already been eroded by recombination in previous generations (see model output, Fig. 2c). Markers that are only weakly linked (e.g. c=0.25) would therefore only be swept in the immediate vicinity of the initial suppressor mutation, and not in the remainder of the species range, diluting the impact seen at these loci.

#### c) Observations of the pattern of genetic variation at suppressor-linked loci post 2006

In order to investigate the longer term impact of the sweep, or to identify its terminus, we also analysed the pattern of variation within the linkage group for a further population sample collected in 2010 (*c.* 32 generations after the 2006 sample). Local LD was still observed, but, as expected from continued recombinational erosion, the extent of LD was reduced (Fig. 6c). In terms of allelic profile, the pattern observed was heterogeneous (Fig. 4; Fig. 5). At six loci (including the three in the direct vicinity of the region containing the suppressor), allele frequencies in the 2010 sample were not significantly different from the 2006 sample. Differentiation of the 2010 to the 2001 samples, as measured by Fst, was equivalent to that previously observed in 2006. The allelic frequency distributions at these loci are at equilibrium, and from these we conclude the sweep had effectively ended. However, allele frequency distributions differed between 2006 and 2010 samples at 5 loci across two genomic regions, one proximal to that carrying the suppressor, and one distal to it. For these, Fst differentiation to the 2001 sample was reduced, and significant differentiation to the 2006 sample was also observed (Fig. 5). In all cases the differentiation is caused by loss (or reduction in frequency) of the major allele that had been swept between 2001 and 2006. Our sampling was geographically broad for both 2006 and 2010 samples, leading us to reject population substructure as a cause of allele frequency heterogeneity.

We finally examined whether the loss of originally swept suppressor-linked alleles by 2010 could be explained by genetic drift. Using the model of Kimura ([17], equation 7) we ascertained the effective population size at which there is a reasonable chance an allele present in 2006 would be absent in 2010. We conducted this for locus G, where the swept allele was at a frequency of 0.22 (n=94) in 2006 and absent in 2010. Setting the time elapsed between samples conservatively at 40 generations (4 years, 10 generations per year based on egg-mature adult period of 36 days), we estimated that loss would occur on 1% of occasions with *N_e_* = 180, and on 5% of occasions with *N_e_*=122. Given the size of Upolu, Samoa, and the ease of collecting adult butterflies (our samples of 48 were collected in two days from three sampling points, and did not involve sampling pupae/larvae), we consider it likely that *N_e_* is considerably larger than this, and thus conclude that drift was unlikely to cause the loss of material observed. Furthermore, we note that loss of introgressed material occurred at two genomic locations that were not in LD with each other, again making chance sampling an unlikely source of the loss of material. Rather, we hypothesize that purifying selection against introgressed material is the most likely reason for the loss of swept alleles.

### The evolution of suppression in Samoa

Our results lead to the following working hypothesis for the evolution of suppression in Samoa. In 2001 the population did not carry the suppressor; nearly all individuals were infected with the male-killer, and the population sex ratio was very female biased, with 100 females per male. Between 2001 and 2005, the suppressor allele was introduced by immigration, consistent with selection occurring in the same genomic region identified as required for male survival in SE Asian butterflies, and the introduction of several novel variants across the swept region. The suppressor was then under very strong selection, as shown by our theoretical model and the rate of change of sex ratio observed in nature [10]: in the first generation after arrival of the immigrant effectively all males in the local mating pool would carry a copy of the suppressor, creating a population frequency of 25%. This intense initial selection was sufficient to produce significant introgression across at least a 25cM region surrounding the locus under selection, which represents the widest genomic region selected during a sweep in a natural population. The sweep had reached equilibrium by 2006. Subsequently, some of the linked genomic regions that had been introduced along with the suppressor declined in frequency, most likely because the introduced alleles were deleterious in the novel Samoan background.

The intensity of selection associated with sex ratio distorting elements may be biologically important because it leads to the invasion of variants that would otherwise be deleterious. The suppressor mutation itself may be highly deleterious in the absence of *Wolbachia* infection, yet invade and become an essential component because it is necessary for male host survival, and its spread does not reduce the frequency of the symbiont (thus it remains required for male survival). It has been widely conjectured that the evolution of sex determination systems might occur in response to the presence of sex ratio distorting microbes, and our data indicates that dramatic changes with associated pleiotropic deleterious effects may be nevertheless be favoured if they rescue males. A pressing area for research is to establish the actual nature of the suppressor mutation (e.g. whether it is part of the sex determination cascade), whether there is a cost of carrying a suppressor in the absence of its benefit from rescuing males, and whether any compensatory evolution has occurred where the suppressor has been present for a longer period.

Beyond the locus under selection, linked variants may also be deleterious. This is one explanation for the gain followed by loss of introgressed material. Our hypothesis is that some introgressed alleles were deleterious on the Samoan genetic background, associated with negative epistasis (Dobzhansky-Muller incompatability) generated during isolation on Samoa. Selection against linked deleterious recessive alleles provides an alternative explanation, although the complete loss observed for some loci suggests that selection against the material is maintained when rare, arguing against recessive effects. Whatever the precise cause of the loss of introgressed material, an important conclusion from our data is that the initial strength of selection would be seriously underestimated from genetic variability data obtained after the final phase of the sweep alone (e.g. 2010 data). Thus, for cases of strong selection over a wide region, caution is needed when estimating the strength of selection from post-sweep genetic data alone.

Finally, our data highlight the profound loss of evolutionary capacity associated with isolation. In Samoa, at least 100 years of extreme sex ratio bias was resolved by a migratory origin of the suppressor that likely involved individual movement over very long distances across the Pacific, rather than *in situ* mutation. One question that still requires answer is ‘why 100 years’. This observation should be considered in concert with our wider knowledge of suppressor spread in this system. First, suppression of male-killing was known in the Phillipines in 1890, but was not observed in neighbouring Borneo (despite presence of a high prevalence male-killing *Wolbachia*) until after 1970 [18]. Suppression was also absent in Hong Kong at this time [18], but present in 2005 [9]. Indeed, by 2005, suppression of male-killing was observed widely across SE Asia [9]. Thus, there was a larger ‘front’ of suppressor-carrying populations building in SE Asia, making arrival of a migrant much more likely after 1970. Evolutionary biologists commonly regard island isolation as a driver of novelty, both in terms of phenotypic traits and endemism. Our study makes it clear that small isolated populations may also be constrained in their response to novel evolutionary challenges. This may be particularly likely for host-parasite symbiosis where there are precise molecular interactions determining parasite persistence and virulence, and few feasible mutations available to provide resistance.

## Materials and methods

### Developing a genome-wide marker set for H. bolina

We utilized high-throughput sequencing of the transcriptome of *H. bolina* to obtain coding sequence from multiple loci across the genome. Following total RNA extraction from 10 *H. bolina* pupae, mRNA library construction and sequencing using the Roche 454 sequencing platform (http://www.454.com), 450 bp reads were *de novo* assembled into contigs using the Newbler assembler to create the first set of Expressed Sequence Tags (EST) for *H. bolina*.

In the absence of any annotated genome or transcriptome for *H. bolina*, the moth *Bombyx mori* was used as a proxy reference genome, this being the only available resource for Lepidoptera at the time of the study. There is a high level of synteny of gene location in the Lepidoptera [19] allowing a targeted gene approach, in which several genes could be selected from each chromosome across the genome. Coding sequence of highly conserved genes such as ribosomal proteins and housekeeping genes from *B. mori* were targeted and then retrieved from NCBI (http://www.ncbi.nlm.nih.gov). To determine putative *H. bolina* orthologs a local tBLASTx was then performed against the *H. bolina* EST set. Only genes that returned a single tBLASTx hit were included, reducing the likelihood of the inclusion of paralogs in our marker set. The orthologous *H. bolina* contigs were then translated into amino acid sequences using the ExPASY online tool (http://web.expasy.org/translate), with the sequence lacking mid-sequence stop codons chosen as the most likely translation. In a final test for paralogs, a reciprocal BLAST was performed of coding sequence from the orthologous *H. bolina* contigs as queries against the *B. mori* genome using the INPARANOID8 search tool ([20]; http://inparanoid.sbc.su.se/).

Intronic regions were targeted for marker development, as they are likely to have a higher degree of nucleotide diversity. Again, conservation of synteny in Lepidoptera genome organisation allowed the intron/exon boundaries in *H. bolina* genes to be inferred using the *B. mori* genome. Through tBLASTx analysis of the *B. mori* coding sequence of the targeted gene against the *B. mori* WGS (Whole Genome Shotgun contigs) database in NCBI, exonic regions were identified (as only these regions will align). The translated orthologous *H. bolina* contig and the corresponding *B. mori* amino acid sequence were aligned using ClustalW [21] and the position of the intron/exon boundaries subsequently located.

Once intron/exon boundaries were identified in *B. mori* genes, and extrapolated to the *H. bolina* orthologous sequences, primers were designed for *H. bolina* that spanned introns of size 500-1000bp (*Bombyx* size approximation). This size range was chosen to enable successful amplification of the intronic region during PCR. Marker optimisation was performed using three test *H. bolina* samples and successful PCR products were sequenced using Sanger technology. From this, 46 markers from 23 chromosomes were chosen that gave clean sequences, with a preference for loci that had 5 or more single nucleotide polymorphisms (SNPs).

### Mapping the suppressor linkage group

In order to investigate the genetic architecture of male-killing suppression in *H. bolina* and determine markers in linkage with the suppressor locus we crossed females of a butterfly population (the Philippines) that were *Wolbachia*-infected and homozygous for the male-killing suppressor allele (***<u>SS</u>***) to males from a *Wolbachia*-infected population (Moorea, French Polynesia) that lacked the suppressor (*ss*), to create suppressor-heterozygous *Wolbachia*-infected offspring (***<u>S</u>****s*) (Fig. 1). Recombination does not occur during female meiosis in the Lepidoptera [22], permitting the progeny of ***<u>S</u>****s* females to be used to identify the linkage group (SLG, Suppressor Linkage group) in which the dominant suppressor allele was carried. To this end, ***<u>S</u>****s* females were crossed with *ss* males to produce the female-informative families. For inclusion in the SLG, markers linked to the suppressor locus are characterized by being present in all surviving sons of the ***<u>S</u>****s* heterozygous mother, rather than the 50% expectation from Mendelian segregation with random survival. Initially each marker was sequenced in the F1 parents (***<u>S</u>****s* female x *ss* male). In each case, SNPs were chosen that were heterozygous in the female and homozygous in the male – following the presumed pattern of the suppressor. These same SNPs were then scored in 16 male and 8 female F2 progeny. Once a marker had been found that was present in half of the daughters (following Mendelian inheritance) but all of the sons (for a son to survive it must have at least one copy of the suppressor, and hence linked marker allele), further markers were developed for that same chromosome based on synteny with *B. mori*. A final suite of 12 markers that produced clean sequence and that spanned the suppressor-associated chromosome were developed to form the SLG.

Recombination does occur in male *H. bolina*, and thus crosses of ***<u>S</u>****s* males to *ss* females (the male-informative families) allow a) mapping of genetic markers within a chromosome relative to each other and b) mapping of the suppressor within the linkage group, in terms of a region of the chromosome that is always present in surviving sons. To this end, the 12 linked markers were sequenced in the female F2 (n = 307) from one male informative cross (***<u>S</u>****s* male x *ss* female) and a linkage map created using JoinMap (version 3.0) [23]. To place the suppressor locus within the map F2 males (n = 60) from this cross were analysed using the same 12 markers. Absence of recombinants in a core subset of markers, flanked by markers with an increasing numbers of recombinants, indicated the position of the suppressor locus (Fig. 2).

### Assessing the effect of suppressor spread on genomic variation

A population sample of butterflies from three time points (2001: n=48, 2006: n=48, 2010: n=46) were collected from one region on the Samoan island of Upolu. For each individual, DNA was extracted using the Qiagen DNEasy kit, and the suite of 12 suppressor-linked markers amplified using PCR. Following Sanger sequencing of the amplicons through both strands, the resultant marker sequences were alignment in Codoncode (http://www.codoncode.com/). SNPs present within and between the population samples were then identified and scored for each individual butterfly. Using the SNP data, the alleles present at each marker in each population sample were estimated using the haplotype reconstruction software PHASE (version 2.1 [24,25]) with 1000 iterations, a thinning interval of 100 and 1000 burn-in iterations. Allele frequencies at each marker for each time group could then be calculated and compared. Output was also examined by eye, with alleles identified first where there was no ambiguity (either homozygous, or a single SNP separating into two defined alleles). Thereafter, alleles were assumed identical to those already identified where possible. The low allelic diversity meant this visual analysis produced very similar result to PHASE output, which can thus be considered robust.

Patterns of differentiation were estimated using GENEPOP [26]. First, heterogeneity of allelic frequency distributions between pairs of time points was estimated using a G test based on allelic frequency distribution. Where allele distributions were heterogeneous, we ascertained the allele whose frequency change made the greatest contribution to heterogeneity as that with the largest standardized residual within the heterogeneity test. This allele was then removed (it was an allele increasing in frequency in each case), and the data retested to ascertain if the population samples were then homogeneous, or whether there was evidence for a second allele that changed in frequency (a second allele was identified in three cases). We additionally used Fst standardized population genetic differentiation to quantify the magnitude of change between allelic frequency distributions between the two samples. In each case, the rare individuals where sequence could not be obtained for particular alleles, or not inferred accurately, were coded as missing information.

Nine unlinked markers, each situated on a different chromosome, were also sequenced for the 2001 and 2006 population samples to investigate the degree to which changes were observed in the wider genome and as a control for demographic effects. These were tested for the presence of heterogeneity between time points using a G test based on allelic frequency distributions, and for differentiation using the Fst statistic.

We additionally analysed evidence for alteration in the pattern of linkage disequilibrium, again using GENEPOP. The significance of LD between all possible combinations of loci was tested in the 2001 and 2006 samples separately. We do not report the magnitude of LD, as this is not a standardized measure, being dependent on the allelic frequency distribution at each locus.

## Acknowledgements

We wish to thank Deborah Charlesworth, Gilean McVean, Pascal Campagne and the Evolution and Ecology of Infectious Disease group at the University of Liverpool, UK, for comments on the manuscript.

## Supporting Information Figure Legends

**Figure S1.**
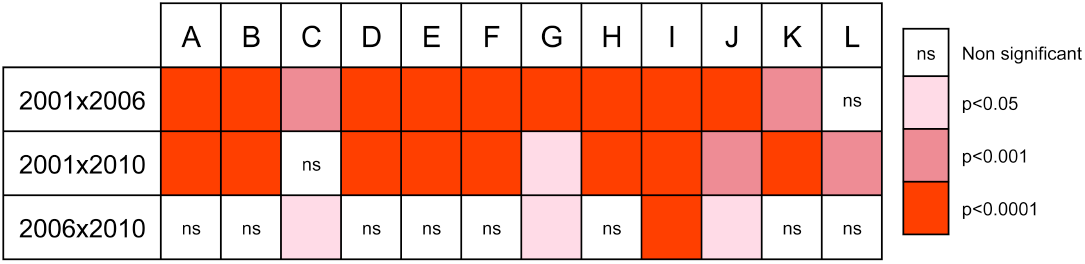
Results of statistical testing for genotypic differentiation between population samples at loci in the linkage group carrying the suppressor. Significance denoted by colour: deep red – significant differentiation as measured at p<0.001; mid red – significant differentiation as measured at p<0.01; pink – significant differentiation as measured at p<0.05, all uncorrected for multiple tests.

**Figure S2.**
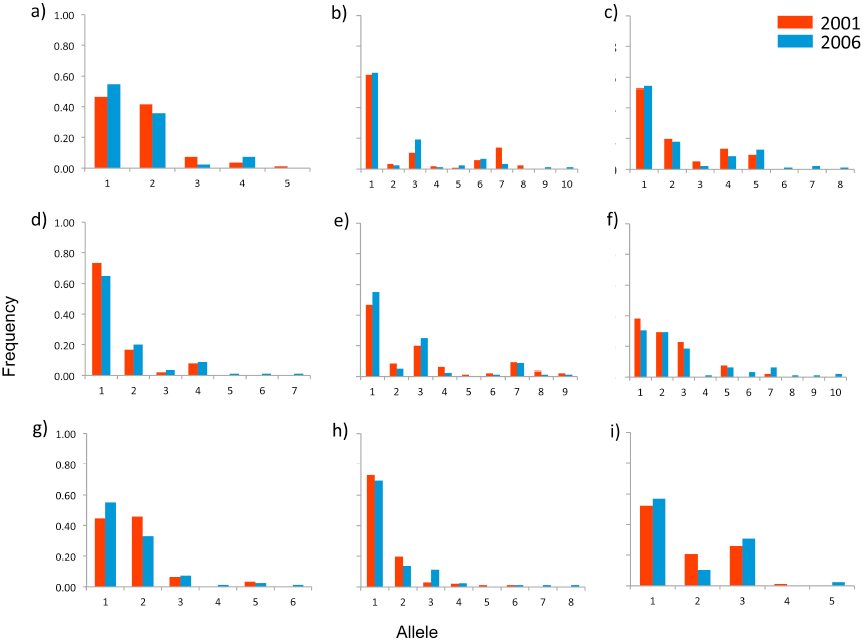
Allelic frequency profiles of markers unlinked to the suppressor. Each of the 9 loci (a to i) is situated within a different chromosome in the *H. bolina* genome, derived from *Bombyx mori* genomic annotations. Allele frequency changes between 2001 (red) and 2006 (blue) at all 9 loci were not significant (Chi Square heterogeneity test; p>0.05, Bonferroni corrected).

**Figure S3.**
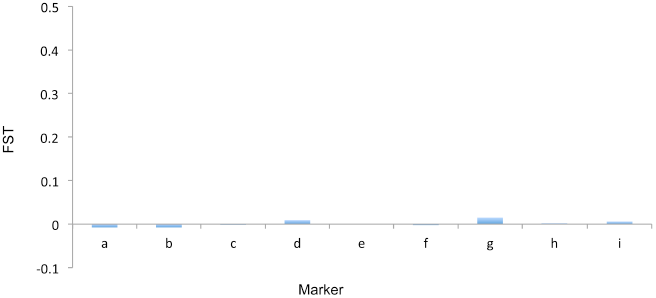
Fst analysis of differentiation between 2001 and 2006 population samples at the nine unlinked loci.

## Text S1: Modelling the spread of the suppressor

### The model

We coded the model in Mathematica.

We model two linked loci, each with two alleles. The first locus can have the wildtype allele *s*, or the male-killing suppression gene *S*. The second locus is linked to the first, and has two selectively neutral alternative alleles denoted *A* and *a*. The model tracks the change in gametic frequencies from one generation to the next. There are four different basic gamete types: *AS*, *As*, *aS*, and *as*. Our individuals are diploid, so these four basic gamete types give nine possible basic individual genotypes: *AASS*, *AaSS*, *aaSS*, *AASs*, *AaSs*, *aaSs*, *AAss*, *Aass*, and *aass*.

The population is infected with *Wolbachia* at a frequency *J*. The infection is vertically transferred from mothers to offspring with 100% efficiency. Thus all of the offspring of an infected mother are infected, and none of the offspring of an uninfected mother are infected. Since there is this difference between males and females, we have to distinguish between male and female gametes. We also have to add infection status to the gametic genotypes to give a total of sixteen gamete types: four basic types, each from a male or female, and either infected or uninfected. Infection status and sex are also added to the individual genotypes to give thirty-six possible genotypes (the nine genotypes from above can be either infected or uninfected, and male or female). Note that for notational ease we still refer to these as “genotypes” although infection status is not a property of the individual’s genome.

The life cycle consists of mating, then selection. The gametic frequencies we begin with give rise to offspring genotypic frequencies. These offspring frequencies undergo selection, and thus result in adult genotypic frequencies. The adults produce gametes, which give us the gametic frequencies for the next generation. Along the way we can record the adult genotypic frequencies and consequently make predictions about the allelic frequencies observed in the real world.

In our model it is not generally the case that there are equal numbers of males and females, because *Wolbachia* affects the two differently. However, there are always equal numbers of male and female gametes (because each mating involves exactly one male and one female). Therefore the female gamete frequencies sum to one, and so do the male gamete frequencies.

Mating is assumed to be at random, and the mating step in our model consists of the transformation of the gametic frequencies into offspring genotypic frequencies. Given a female gamete at frequency *f*, and a male gamete at frequency *m*, the frequency of offspring resulting from combining these two gamete types will be the product of their frequencies *fm*. An added complication arises due to cytoplasmic incompatibility (CI). This is when mating between infected males and uninfected females result in fewer offspring than other matings. For simplicity, we assume that CI is total, so that fusions between infected male gametes and uninfected female gametes result in no offspring. We then renormalise the offspring frequencies appropriately so that they still sum to one for both males and females.

Once we have the offspring genotypes, selection occurs, transforming the offspring genotypic frequencies into adult genotypic frequencies. In our model, the only element of selection is male-killing: any infected males lacking the *S* suppressor gene (i.e. those with genotype *ss*) are killed by the *Wolbachia*. Males heterozygous at the suppressor locus (i.e. with the genotype *Ss*) are killed by their infection 50% of the time. Males homozygous for the suppressor (i.e. those with genotype *SS*) are not killed. For simplicity we assume no other selective effects. Notably this means that we are modelling the situation in which neither the suppressor nor the linked allele impose costs on their bearers.

Because females are unaffected by *Wolbachia*, there are now more females than males in our population. This will affect the observed allelic frequencies, since the allelic frequencies in males and females will differ (*s* genes, for example, will be more common in females because they are selectively neutral to a female, while males bearing *s* genes are more likely to die through male-killing). To account for this fact we renormalise the post-selection genotypic frequencies so that the sum of both male and female genotypic frequencies is one. This gives us the adult genotypic frequencies. From this data we get the model’s predictions for observed frequency of *A* and *S* alleles.

To complete the generation, we finally transform the adult genotypic frequencies into gametic frequencies. This is trivial in the case of most of the genotypes. However, for genotypes *AaSs*, things are more complicated. This is because *AaSs* individuals could have been formed by *AS* x *as* crosses, or by *As* x *aS* crosses. Denote the probability that a randomly-chosen *AaSs* individual was formed by an *AS* x *as* cross by *μ*, and the probability of recombination between the two loci of interest by *r*. Then the frequency of gamete types from infected *AaSs* individuals is

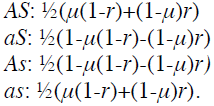

If we denote the probability that a randomly-chosen uninfected *AaSs* individual was formed by an *AS* x *as* cross by *ν* we can produce similar frequencies. It remains only to find *μ* and *ν*. But since we know the previous generation’s gamete frequencies this is trivial.

In the initial population of size *N* we suppose that the *A* and *S* alleles are absent. With a proportion *J* of the population infected with *Wolbachia*, the gamete types and frequencies in the population are therefore infected female *as* (frequency *J*), uninfected female *as* (frequency 1 – *J*), and uninfected male *as* (frequency 1). At this point the sex ratio is (1 - *J*)/(2 - *J*). We then introduce an immigrant of known sex and genotype into the population. To do this, we multiply the male gamete frequencies by *N*(1 - *J*)/(2 - *J*), and the female gamete frequencies by *N*/(2 - *J*) to give a “gamete mass” measurement. Then, given the immigrant’s genotype we get the probability that it produces a gamete of each type (we assume for simplicity that *μ* = *ν* = ¼). We then add the immigrant gamete probabilities to the gamete mass measurement, and renormalise so that female gamete frequencies sum to one, and so do male gamete frequencies. This gives us the initial gamete frequencies. As an example, in the case of a single infected male immigrant of genotype *AASS*, the initial gametic frequencies are:

Infected female *AS*: 0

Infected female *aS*: 0

Infected female *As*: 0

Infected female *as*: *J*

Uninfected female *AS*: 0

Uninfected female *aS*: 0

Uninfected female *As*: 0

Uninfected female *as*: 1 – *J*

Infected male *AS*: (2 - *J*)/(2 – *J* + (1 - *J*)*N*)

Infected male *aS*: 0

Infected male *As*: 0

Infected male *as*: 0

Uninfected male *AS*: 0

Uninfected male *aS*: 0

Uninfected male *As*: 0

Uninfected male *as*: (1 - *J*)*N*/(2 – *J* + (1 - *J*)*N*)

### Local LD

Modelling local linkage disequilibrium is similar to above, with the added complexity that there are now two neutral loci to keep track of rather than just one. We label the alleles loci *A*/*a*, *B*/*b*, and *S*/*s* (the suppressor). Thus there are now 27 basic genotypes (3^3, since there are 3 loci, and three different possible genotypes at each one).

We now have to separately consider recombination between the suppressor and the two neutral loci (occurs at a rate *r*_1_), and recombination the two neutral loci (occurs at a rate *r*_2_). We make the simplifying assumption that these recombination events occur independently of one another, which is probably a reasonable assumption as long as the two rates aren’t too similar in size.

The final complication is that with so many more genotypes to keep track of, there are many more types of heterozygotes. In line with our calculation of *μ* and *ν* above, we have to calculate the frequency of each type of gamete being passed on by a given heterozygote.

